# Overexpression of enhanced yellow fluorescent protein fused with Channelrhodopsin-2 causes contractile dysfunction in skeletal muscle

**DOI:** 10.1101/2024.06.06.597782

**Authors:** Syeda N. Lamia, Carol S. Davis, Peter C.D. Macpherson, T. Brad Willingham, Yingfan Zhang, Chengyu Liu, Leanne Iannucci, Elahe Ganji, Desmond Harden, Iman Bhattacharya, Adam C. Abraham, Susan V. Brooks, Brian Glancy, Megan L. Killian

## Abstract

Skeletal muscle activation using optogenetics has emerged as a promising technique for inducing noninvasive muscle contraction and assessing muscle function both in vivo and in vitro. Transgenic mice overexpressing the optogenetic fusion protein, Channelphodopsin2-EYFP (ChR2-EYFP) in skeletal muscle are widely used; however, overexpression of fluorescent proteins can negatively impact the functionality of activable tissues. In this study, we characterized the contractile properties of ChR2-EYFP skeletal muscle and introduced the ChR2-only mouse model that expresses light-responsive ChR2 without the fluorescent EYFP in their skeletal muscles. We found a significant reduction in the contractile ability of ChR2-EYFP muscles compared to ChR2-only and WT mice, observed under both electrical and optogenetic stimulation paradigms. Bulk RNAseq identified downregulation of genes associated with transmembrane transport and metabolism in ChR2-EYFP muscle, while the ChR2-only muscle did not demonstrate any notable deviations from WT muscle. The RNAseq results were further corroborated by a reduced protein-level expression of ion-channel-related HCN2 in ChR2-EYFP muscles and gluconeogenesis-modulating FBP2 in both ChR2-EYFP and ChR2-only muscles. Overall, this study reveals an intrinsic skeletal dysfunction in the widely used ChR2-EYFP mice model and underscores the importance of considering alternative optogenetic models, such as the ChR2-only, for future research in skeletal muscle optogenetics.

## Introduction

The ability to control skeletal muscle activation is essential for assessing muscle function both in vivo and in vitro. In a controlled laboratory environment, skeletal muscle activation is predominantly assessed using either nerve-mediated or direct electrical stimulation. While electrical stimulation techniques are well established and repeatable, significant technical challenges exist, including the iatrogenic effects of needle insertions and the inability to activate denervated muscles, which is common in peripheral nerve injuries (e.g., brachial plexus avulsion and spinal cord injuries) and neuromuscular disorders (e.g., muscular dystrophies and amyotrophic lateral sclerosis) (*1–3*). Furthermore, the use of established electrical stimulation techniques presents significant challenges when applied to neonatal mice and preclinical models used for studying muscle development and pathology. Smaller muscles in neonatal mice and less resilient muscle in pathological mouse models can exacerbate iatrogenic issues due to repeated needle insertions. Consequently, traditional electrical stimulation methods might compromise the integrity of muscle structure, affect perfusion, and induce inflammation, thereby diminishing the physiological relevance of experimental outcomes.

To overcome these challenges, skeletal muscle activation with optogenetic stimuli (i.e., optogenetics) has emerged as a noninvasive technique to induce muscle contraction (*4–8*). Optogenetic techniques are well established in neurobiological research due to high spatio-temporal resolution (*9*, *10*). Optogenetic stimulation can elicit muscle contraction via depolarization of muscle fibers that express light sensitive opsins, such as Channelrhodopsin-2 (ChR2) protein (*4*, *5*). ChR2 is a nonselective transmembrane cation channel derived from the green microalgae *Chlamydomonas reinhardtii* (*9*). When exposed to blue light (455 nm), the transmembrane ChR2 opens to allow non-selective cation flow for muscle fiber depolarization and contraction (*10*, *11*). Thus, optogenetic contractions can be elicited independent of motoneuron excitation, leading to muscle stimulation even when muscle is denervated. Direct optogenetic stimulation of excitable muscle fibers have been used to induce contraction noninvasively via transdermal light exposure (*1*, *4–7*) and serve as an alternative to electrical stimulation (*1*). Optogenetic stimulation has primarily been conducted using transgenic mice expressing ChR2 (H134R)-enhanced yellow fluorescent protein (EYFP) fusion (commonly known as Ai32) under the Sim1 and chicken-β-actin promoters (*4*, *5*). In these mice, skeletal muscle localizes ChR2-EYFP within the sarcolemma and t-tubules, as well as within voltage gated sodium channels of the sarcolemma (*4*). Our group has previously used the established Ai32 transgenic strain to achieve optogenetic control of skeletal muscle using the Acta1 promoter, and we showed that ChR2-EYFP was expressed only in the skeletal muscle fibers and not in the nerves (*6*).

While the EYFP present in the commonly used Ai32 mouse line is useful for detecting Cre recombination within tissues, fluorescent reporters such as EYFP and green fluorescent protein (GFP) have been associated with cytotoxicity (*12–15*). Moreover, Agbulut et al. demonstrated that cytosolic GFP in muscle impairs actin-myosin head binding, leading to contractile dysfunction (*16*). High levels of ChR2-EYFP expression have also been shown to alter the electrophysiological properties and morphology of neurons (*17–19*). Furthermore, ChR2-EYFP mediated immunogenicity has been reported to cause motor neuron death and muscle atrophy (*20*). We have previously shown that the doxycycline-inducible Acta1-tetO-rtTA-Cre; Ai32 mouse model exhibited aggregation of EYFP in skeletal muscle (*6*). We were concerned that this clustering of EYFP had the potential to influence contractile and electrophysiological function of muscle. To address this concern, we deleted the EYFP gene from the ChR2-EYFP mice using CRISPR/Cas9. To test if and how EYFP influences skeletal muscle contractility, we compared contractile properties using electrical stimulation as well as optogenetics using multiscale approaches. We also evaluated the effect of EYFP on light-induced contractile force in vitro with collimated blue light. In addition, we determined how the presence of ChR2/EYFP affects muscle ultrastructure and fiber size. To obtain a mechanistic view behind the contractile dysfunction, we used RNA sequencing (RNAseq) to identify the transcriptional response of muscle following daily bouts of optogenetic stimulation in these two ChR2-expressing optogenetic strains.

## Results

### High ChR2-EYFP expression and vacuole formation observed in ChR2-EYFP muscles

To investigate the effects of ChR2-EYFP overexpression in skeletal muscle, we generated a transgenic mouse line to express ChR2 in skeletal muscle with EYFP fluorescent reporter (*6*, *7*). Doxycycline-treated, ACTA1-rtTA;tetO-Cre mice were bred with ChR2-EYFP^fl/fl^ mice (known as Ai32 mice) to generate Acta1Cre;ChR2-EYFP^fl/fl^ mice, referred to as ChR2-EYFP mice onward. We observed potentially abnormal expression of ChR2-EYFP fusion protein in the gastrocnemius muscle sarcolemma of 2-month-old ChR2-EYFP mouse (Fig. 2A). To determine if this phenomenon was independent of Cre expression, we visualized EYFP expression in other muscle-specific Cre-lines, such as CKCre;ChR2-EYFP^fl/fl^ (Fig. 2A), Myf6Cre;ChR2-EYFP^fl/fl^, and Sim1Cre;ChR2-EYFP^fl/fl^ (supplementary Fig. S2). In all cases we observed overexpression of ChR2-EYFP protein in the sarcolemma. We labeled ChR2-only muscle with cell membrane dye Di-2-ANEPEQ which demonstrated regular muscle structure (supplementary Fig. S2). Using transmission electron microscopy (TEM), we identified accumulation of vacuoles in the ChR2-EYFP EDL muscle along the sarcolemma (Fig. 2B) which are not seen in Wildtype (WT) muscles. Moreover, the overexpression induced distinct macro-morphological alterations in muscle tissues. Specifically, the overexpressed protein resulted in the formation of voids between parallel muscle cells and protrusions within the muscle cells.

**Fig. 1.**
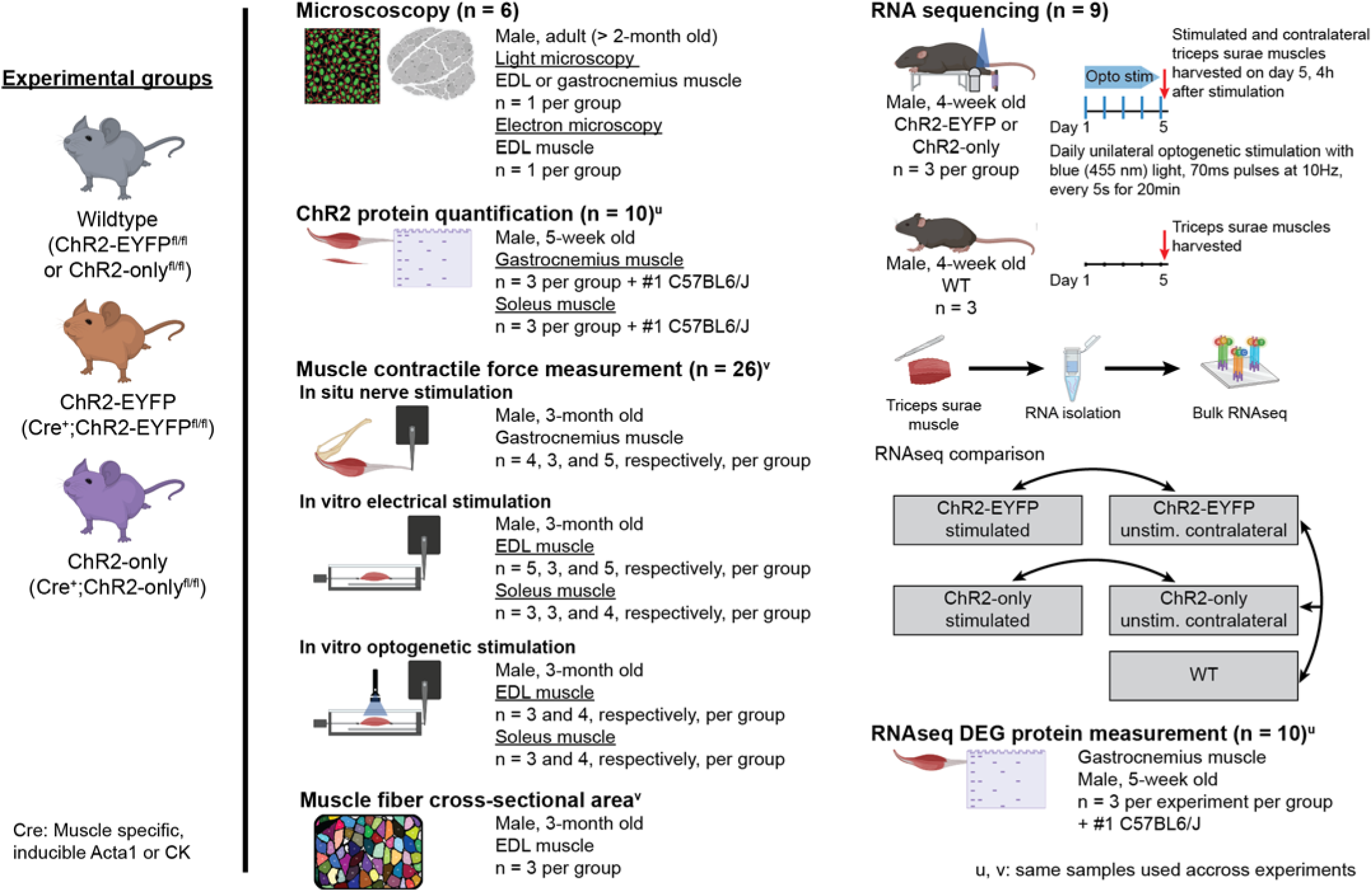
Study design: schematic of experiments with mouse strains and number of mice utilized. Mice with muscle specific expression of optogenetic ChR2-EYFP fusion or ChR2-only protein (referred to as ChR2-EYFP or ChR2-only strains, respectively) and their wildtype littermates were used in various experiments. Young adult (2-3-month-old) mice were used in microscopy, muscle contractility with both optogenetic and gold standard nerve or electrical stimulation, and fiber cross sectional area measurement. Muscles from young (5-week-old) mice were harvested for protein measurements. For RNA sequencing, right triceps surae muscles of young (4-week-old) ChR2-EYFP and ChR2-only mice were stimulated with blue light (455 nm) to induce optogenetic muscle contraction. After 5 daily bouts of 20 min stimulation, stimulated and age matched wildtype mice were euthanized to harvest the triceps surae for sequencing. The superscripts ‘u’ or ‘v’ indicate same mice were used across different experiments. The number of mice used in each experiment are presented as WT, ChR2-EYFP, and ChR2-only, respectively.

**Fig. 2.**
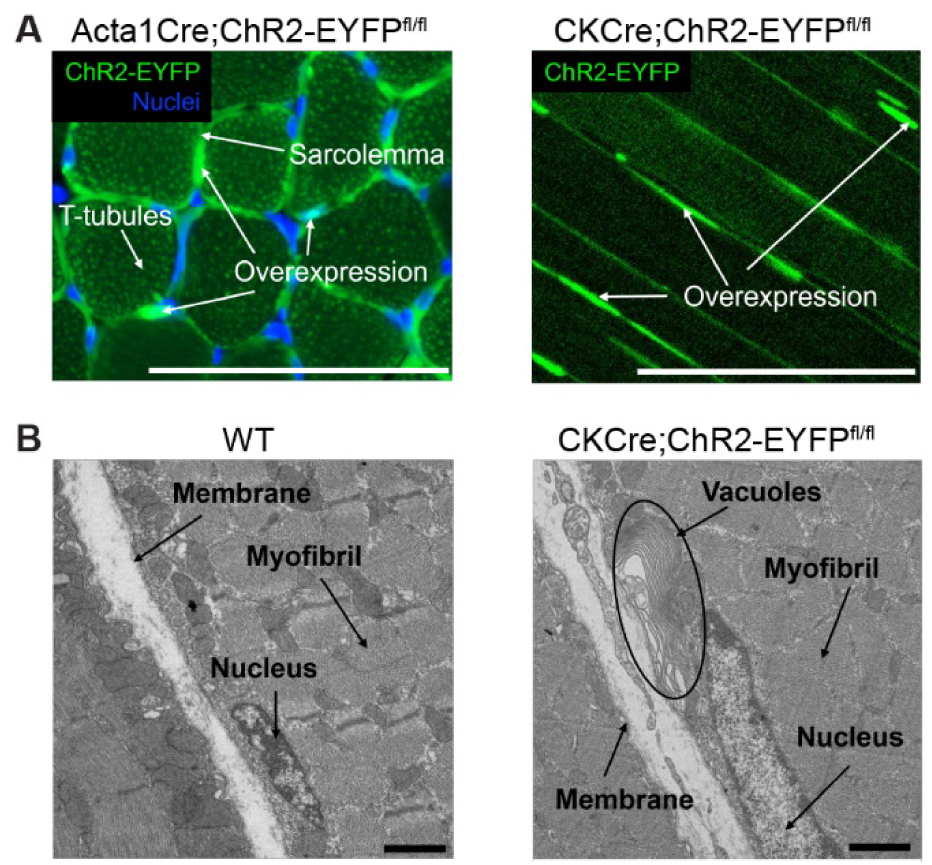
Abnormal ChR2-EYFP clustering and vacuole formation was observed in optogenetic mouse skeletal muscle. (A) Confocal imaging demonstrated ChR2-EYFP overexpression in the transverse plane of gastrocnemius muscle of inducible Acta1Cre;ChR2-EYFP^fl/fl^ mouse and the longitudinal plane of the EDL muscle of CKCre;ChR2-EYFP^fl/fl^ mouse. (B) Transmission electron microscopy of WT EDL muscle and CKCre;ChR2-EYFP EDL muscle revealed the presence of abnormal vacuoles (circled) with the expression of ChR2-EYFP. n = 1 per group. Scale bar: A and B, 100 μm; C and D, 1 μm.

### ChR2 protein expression was reduced in ChR2-only compared to ChR2-EYFP gastrocnemius muscles

Next, we bred doxycycline-treated, ACTA1-rtTA;tetO-Cre mice with ChR2-only^fl/fl^ mice (developed in Glancy lab, see methods for details) to generate Acta1Cre;ChR2-only^fl/fl^ mice, referred to as ChR2-only mice onward. We measured ChR2 protein levels in the muscles of the 5-week-old ChR2-EYFP and ChR2-only mice to determine if ChR2 expression is different between the strains. ChR2 protein was identified at two different molecular weights (55 kDa and 35 kDa, respectively) in ChR2-EYFP and ChR2-only groups (Fig. 3). The ChR2-EYFP gastrocnemius muscles had significantly higher expression of ChR2 protein than the ChR2-only gastrocnemius muscles (Fig. 3.3A), however, no difference in ChR2 expression was observed in the soleus muscle (Fig. 3.B). The larger molecular weight for ChR2-EYFP was expected given the fusion with EYFP (∼25 kDa).

**Fig. 3.**
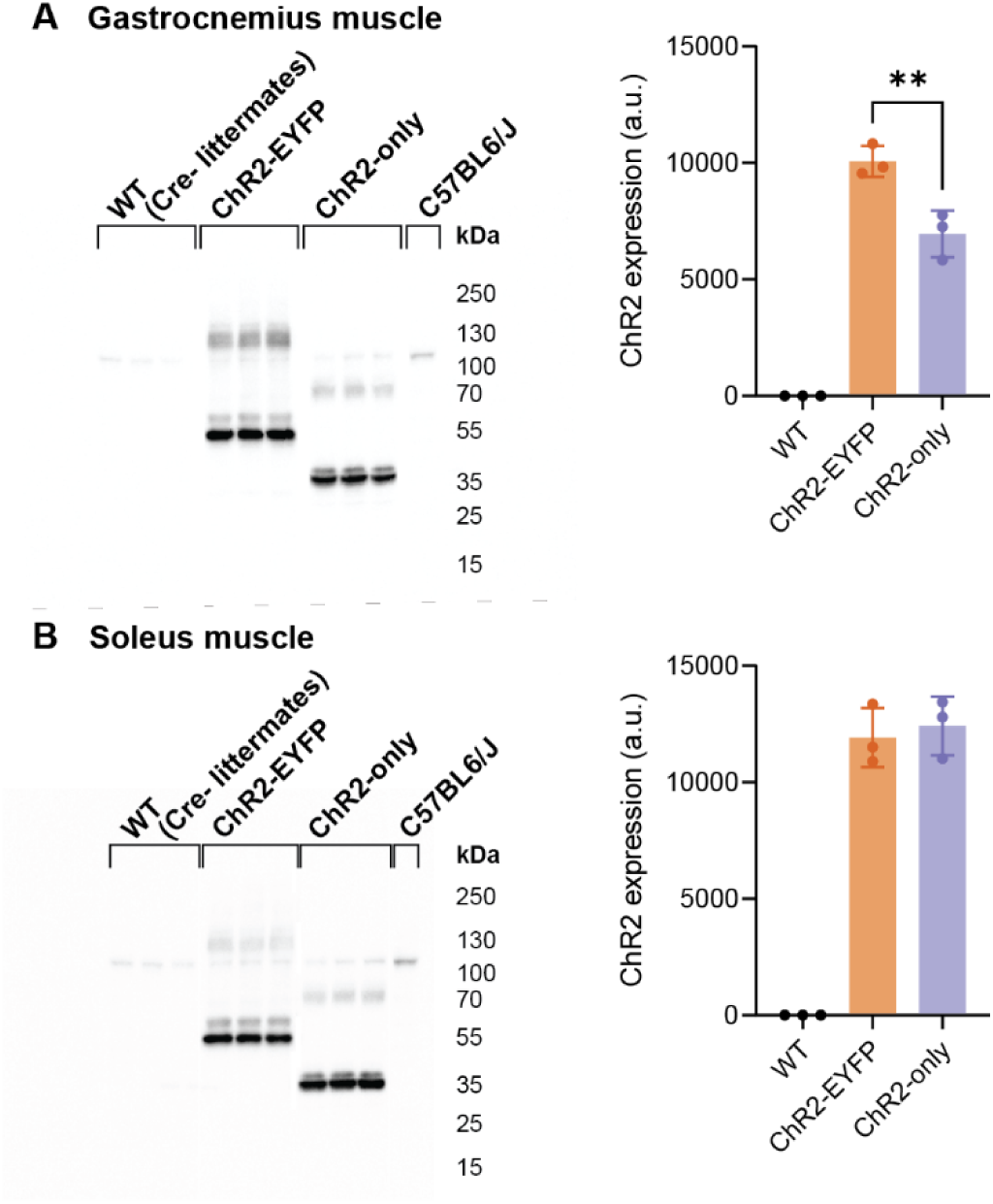
ChR2-EYFP gastrocnemius muscle expressed significantly higher ChR2 protein than the ChR2-only gastrocnemius muscle, but no difference was observed in the soleus muscle. ChR2 western blot and quantification of (A) gastrocnemius and (B) soleus muscle. n = 3 per group. Data were compared using one-way analysis of variance (ANOVA) with Sidak’s correction for multiple comparisons. Error bars denote means ± SD. ** p<0.01.

### The presence of ChR2-EYFP fusion protein led to force deficits in fast-twitch muscle

We found that the 3-month-old ChR2-EYFP mice were smaller than age-matched WT and ChR2-only mice (Fig. 4A). To understand if and how the ChR2-EYFP fusion protein was influencing the growth and function of skeletal muscle, we evaluated the individual weights and force generation potential of muscles with a battery of *in situ* and *in vitro* tests. The mass of gastrocnemius muscles from ChR2-EYFP mice were significantly lower than age- and sex-matched WT or ChR2-only strains (Fig. 4B). Isometric tetanic force of ChR2-EYFP gastrocnemius muscles were also significantly reduced compared with WT and ChR2-only mice at all test frequencies except for 40 Hz (Fig. 4C). To account for differences in mouse and muscle size, we normalized the maximum tetanic force with respect to muscle’s physiological cross-sectional area (PCSA) to obtain the maximum specific tetanic force (Equation 1). However, the normalized force remained significantly lower in the ChR2-EYFP gastrocnemius muscle compared with WT and ChR2-only muscles (Fig. 4D), suggesting the functional deficit was not scaled to animal/muscle size alone.

**Fig. 4.**
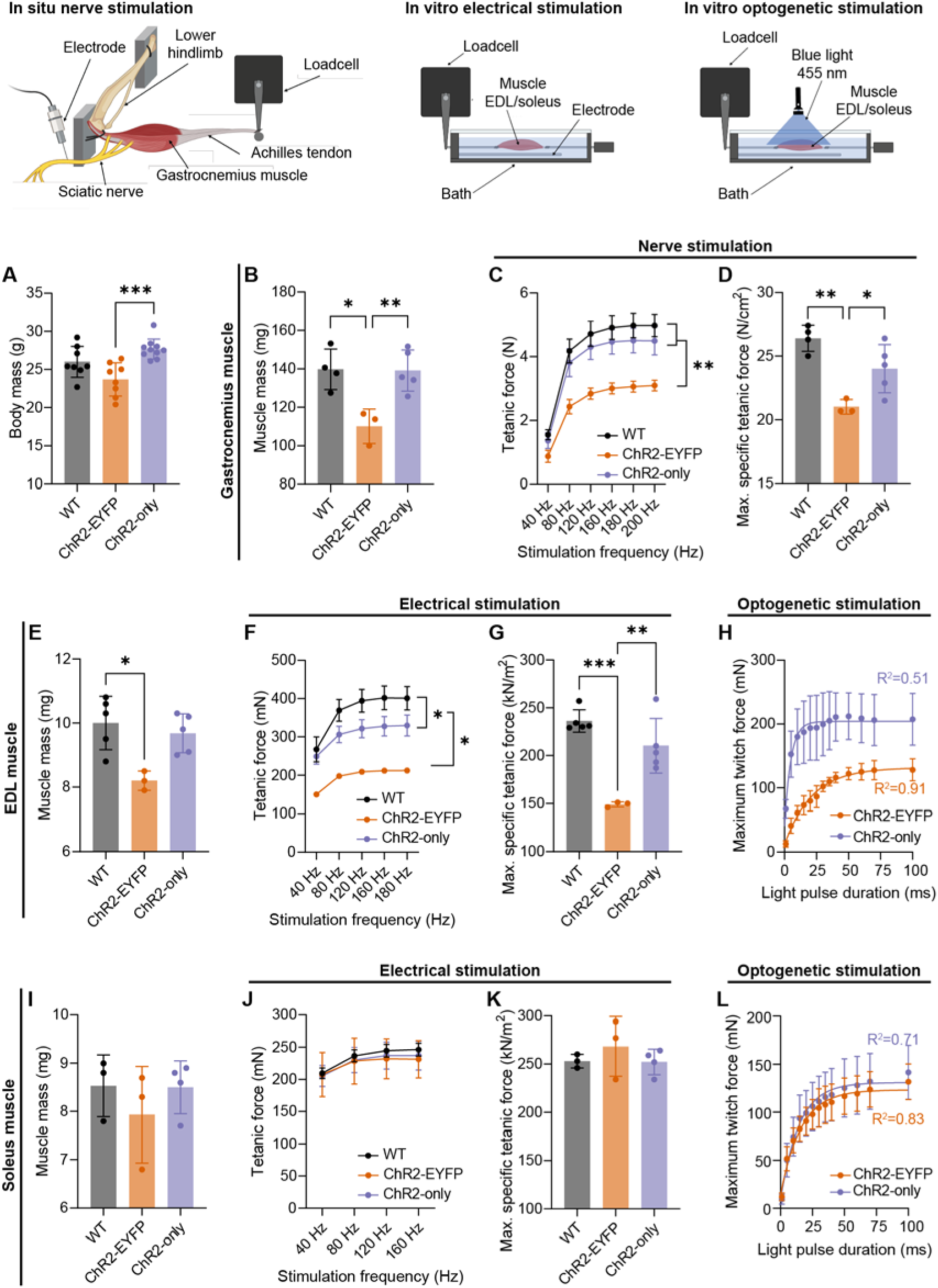
The presence of EYFP with ChR2 compromised the electrical and optogenetic contractile properties of fast-twitch skeletal muscle compared to ChR2 only and WT muscle. (**A**) Body mass of all mice used in the experiment (N = 26; n≥8 per genotype). (**B, E, and I**) Gastrocnemius and EDL, but not soleus muscle masses, were significantly different in ChR2-EYFP mice compared to WT and ChR2-only mice. (**C, F, and J**) Tetanic forces of the gastrocnemius and EDL, but not soleus muscles, were significantly reduced in ChR2-EYFP mice compared to WT and ChR2-only at incremental electrical stimulation frequencies. (**D, G, and K**) Maximum specific tetanic forces of the gastrocnemius, EDL, and soleus muscles, respectively; and (**H and L**) Light induced twitch forces of EDL and soleus muscle, respectively, measured at light pulse durations ranging from 1-100 ms. For (A), (B), (D), (E), (G), (I), and (K), data were compared using one-way ANOVA with Tukey’s correction for multiple comparisons. For (C), (F), and (J), data were compared using repeated measures two-way ANOVA with Tukey’s correction. For (H) and (L), data points from the average maximum twitch within strains were fitted into one-phase decay model. Error bars denote means ± SD. * p<0.05, ** p<0.01, and *** p<0.001. EDL, extensor digitorum longus. Each dot represents a biological replicate (individual mouse).

Next, we evaluated direct muscle contractility *in vitro* to assess muscle function independent of motor neuron innervation. The EDL, a muscle composed primarily of fast-twitch, glycolytic fibers, also weighed significantly less in ChR2-EYFP mice compared to EDLs of WT mice (Fig. 4E). Tetanic forces were significantly different between the three groups (Fig. 4F). The normalized maximum specific tetanic force of the EDL muscle was significantly lower in ChR2-EYFP EDLs compared with WT and ChR2-only EDLs, however no difference was observed in ChR2-only and WT muscles (Tukey’s multiple comparisons adjusted p= 0.14, Fig. 4G). This deficit in direct muscle contractility in the ChR2-EYFP EDLs was further tested in our optogenetic strains using blue light activation. ChR2-only EDL muscles produced higher twitch forces than ChR2-EYFP muscle at all pulse duration (Fig. 4H). We modeled the force response to pulse duration using Akike information criteria (AIC) using one phase decay (R^2^=0.51 and 0.91, respectively, for ChR2-only and ChR2-EYFP) and measured higher contractile forces at smaller pulses in ChR2-only EDLs compared to ChR2-EYFP EDLs. Specifically, ChR2-only EDL muscles twitch force plateaued at shorter pulse durations (i.e., 30 ms duration) compared to ChR2-EYFP EDL muscles (100 ms). Similarly, ChR2-only EDLs generated higher forces with partially fused tetanus at 50 Hz, 10 ms pulse (supplementary Fig. S3), which was not achieved in ChR2-EYFP EDLs.

To determine if this force deficit was dependent on fiber types, we measured the contractile properties of the soleus muscle, which comprise a substantial population of slow-twitch, oxidative fibers, using *in vitro* contractility experiments. Contrary to what we found in fast-twitch muscles, we did not observe any differences in muscle mass (Fig. 4I), isometric tetanic force (Fig. 4J), or maximum specific tetanic force (Fig. 4K), suggesting that the contributions of EYFP in contractility in muscle-specific optogenetic strains may be fiber-type dependent. The optogenetic twitch forces in soleus muscle was also similar in both strains at all pulse durations and they plateaued at similar pulse durations (70 ms, Fig. 4L, AIC-validated one phase decay, R^2^> 0.7 for both strains). Soleus muscles from ChR2-EYFP and ChR2-only generated similar forces with partially fused tetanus at 50 Hz, 10 ms pulse (supplementary Fig. S3).

To investigate if changes in calcium handling mechanism of muscle were influenced by EYFP expression, we recorded the twitch times of EDL and soleus muscles during *in vitro* experiments. We observed a small but significant increase in the half-relaxation times of EDL and soleus muscles (supplementary Fig. S3) suggesting potential alterations in calcium machineries in muscle fibers.

### ChR2-EYFP EDL muscle fibers were smaller compared to WT and ChR2-only fibers

Because we observed a significant decrease in ChR2-EYFP EDL muscle mass compared to WT and lower specific force production of ChR2-EYFP muscles, we next determined the muscle fiber sizes in these strains histologically at 3-month age (Fig. 5A). Muscle fiber area of both optogenetic strains were smaller compared to WT muscle, with ChR2-EYFP displaying a significantly reduced average fiber size (Fig. 5B). While WT muscle demonstrated a uniform distribution of fiber CSA, ChR2-EYFP muscles had higher percentage of smaller fibers (Fig. 5C).

**Fig. 5.**
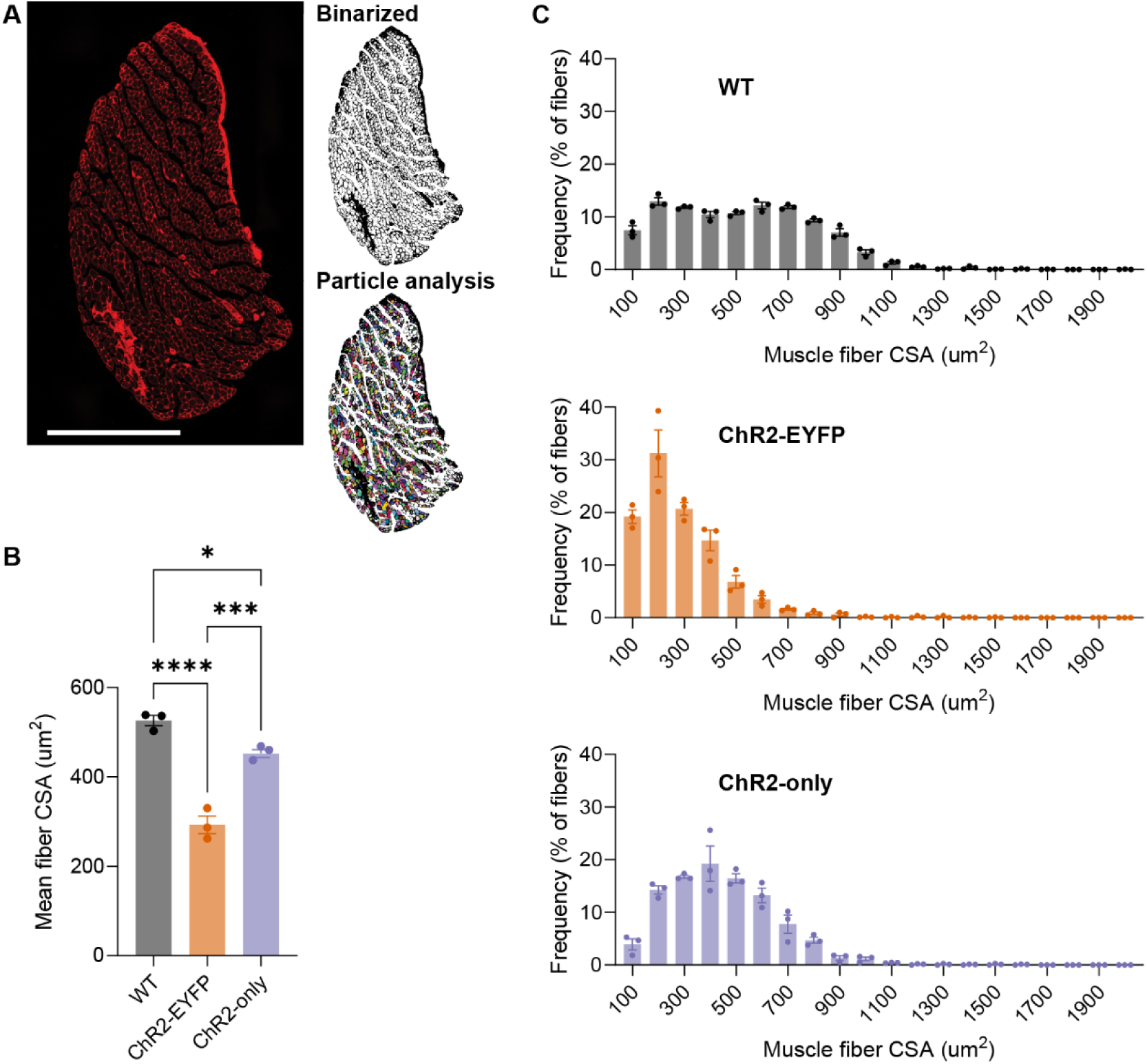
The presence of EYFP in ChR2-expressing EDL muscle led to smaller fiber size. (A) WGA-stained EDL muscle section. Images were binarized before particle analysis. (B) Mean muscle fiber area; and (C) fiber size frequency distribution. WGA, wheat germ agglutinin; CSA, cross-sectional area. Scale bar = 500 μm. n =3 per group. Error bars denote mean±SEM. * p<0.05, *** p<0.001, and **** p<0.0001. For (B), data were compared using one-way ANOVA with Tukey’s multiple correction.

### Unstimulated ChR2-EYFP muscle were transcriptionally unique compared to ChR2-only and WT muscle

To evaluate if the muscles from the ChR2-EYFP and ChR2-only optogenetic strains respond differently to daily repeated optogenetic stimulation, we unilaterally stimulated the triceps surae muscles of 4-week-old mice from these two strains for 5 days (Fig. 1). In total, we ran four RNAseq comparisons as shown in Supplementary Table S1. Transcriptomes of WT and unstimulated ChR2-only muscles were in general similar, as shown by the unsupervised clustering of WT and unstimulated ChR2-only samples in the principal component analysis (Fig. 6A). We identified 1,283 genes that were differentially expressed in the ChR2-EYFP and 410 in the ChR2-only muscle compared to WT, respectively (Table 1). Among the 632 and 251 downregulated genes in ChR2-EYFP and ChR2-only muscles, respectively, 142 were common in both strains. We identified 44 overlapping upregulated genes between ChR2-EYFP and ChR2-only muscles. We used the Database for annotation, visualization, and integrated discovery (DAVID) and found that downregulated genes in ChR2-EYFP muscle transcriptome enriched metabolism, phosphorylation, transmembrane transport, and muscle contraction biological processes (Fig. 6B). Upregulated genes in ChR2-EYFP muscle were associated with inflammatory response and extracellular matrix (ECM) organization (Fig. 6C). To understand the magnitude of differential expression between genotypes at baseline (unstimulated), we plotted the log_2_(FoldChanges; FC) of genes associated with several processes such as muscle contraction, gluconeogenesis, collagen accumulation and degradation, and transmembrane (ion channel) transport (Fig. 6D-G, respectively). We observed downregulation in fast-twitch muscle contraction related genes such as *fMybpc2, Mylpf, Tnni2, and Tnnt3,* which may explain the loss of force in fast-twitch EDL muscle (*21–24*) but not in the slow-twitch soleus muscle. In ChR2-EYFP muscle, we observed a downregulation of transmembrane transport genes that encode the voltage-gated potassium and sodium ion channels (*Hcn2, Kcnf1, Kcnc3, Kcnb1*, etc.) compared to WT muscle. While ECM degradation-associated genes such as *Mmps*, *Ctss*, and *Ctsk* were mostly upregulated in the ChR2-EYFP muscle, MMP inhibitor *Timp1* expression was also upregulated. Genes modulating gluconeogenesis such as *Fbp2*, *Pgam2*, and *Gpd1* were downregulated in ChR2-EYFP muscle, however, they were unchanged in the ChR2-only muscle. To assess if reduced gene expression was translated to protein level, we measured expression of several targets: fructose-1,6-bisphosphatase 2 (FBP2), actinin alpha 3 (ACTN3), tissue inhibitor of metalloproteinases 1 (TIMP1), and hyperpolarization activated cyclic nucleotide gated potassium channel 2 (HCN2) in 5-week-old gastrocnemius muscles. Both ChR2-EYFP and ChR2-only muscles showed decreased FBP2 protein expression compared to WT (Fig. 7A). HCN2 protein was significantly reduced at the protein level in ChR2-EYFP gastrocnemius muscle compared to ChR2-only muscle (Fig. 7B). However, ACTN3 and TIMP1 protein expression did not match the transcription levels (Fig. 7, and supplementary Fig. S5). No biological processes were significantly enriched by the ChR2-only DEGs compared to the WT muscle transcriptome (false discovery rate (FDR) < 0.1).

**Fig. 6.**
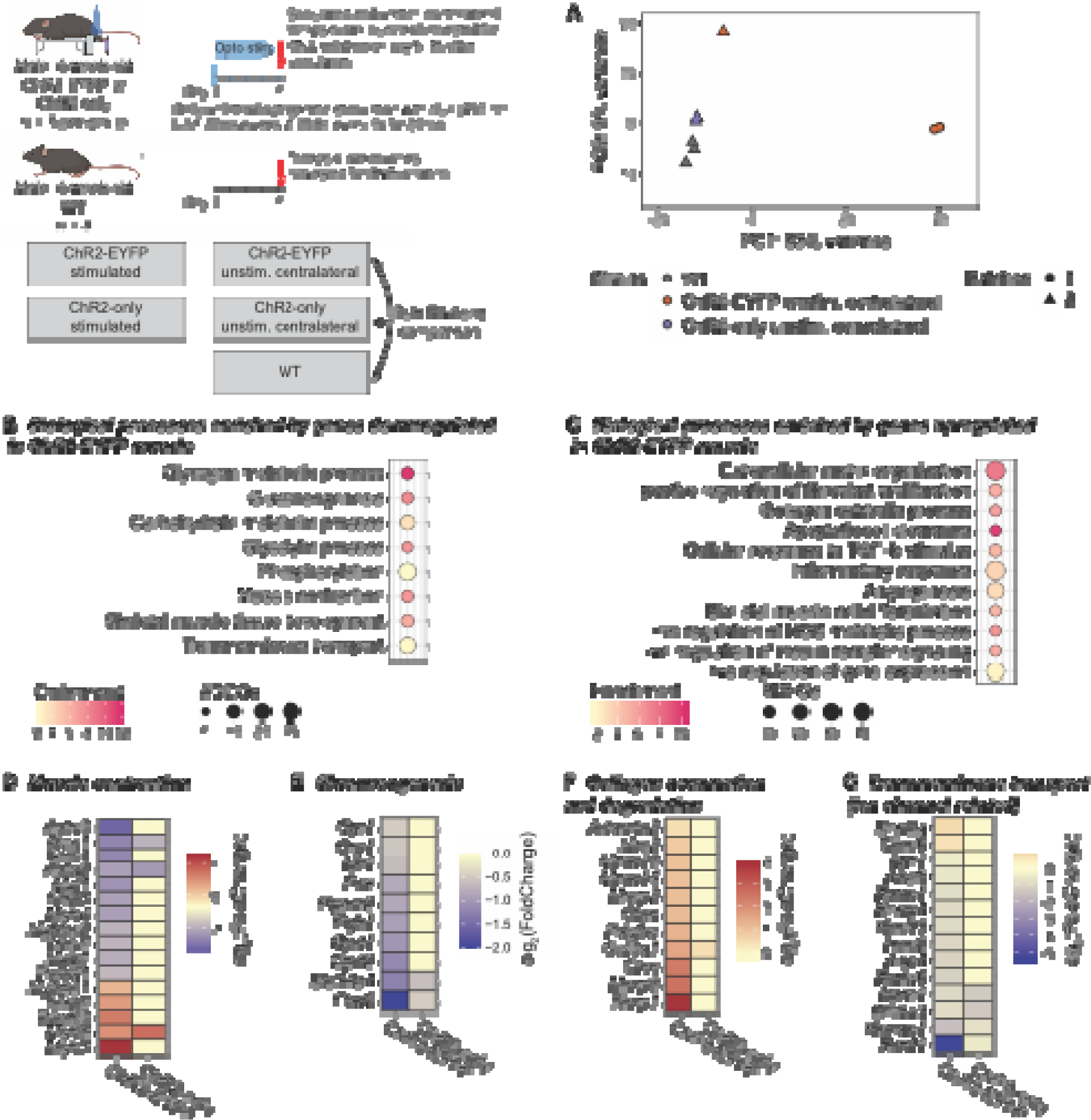
RNA sequencing revealed differential gene expression between WT, unstimulated ChR2-EYFP, and unstimulated ChR2-only triceps surae (i.e., gastrocnemius and soleus) muscles. (A) Principal component analysis of gene expression shows clustering of WT and ChR2-only muscles. Shapes indicate different sequencing batches. Biological processes enriched in ChR2-EYFP skeletal muscle compared to WT indicated (B) downregulation of metabolism, muscle contraction, and transmembrane transport related genes and (C) upregulation of inflammatory response and ECM related genes. (D-F) Gene expressions associated with enriched muscle contraction, gluconeogenesis, and collagen catabolism biological processes, respectively. n = 3 per group.

**Table 1.** Percentages of total light absorbed within volume or lost at each boundary.

|  | % Light<br>Absorbed within<br>Boundaries | % Light<br>Reflected | % Light<br>Lost to<br>Edges | % Light<br>Transmitted |
| --- | --- | --- | --- | --- |
| in vitro | 23.13 | 19.07 | 54.86 | 2.94 |
| In vivo | 37.6 | 30.40 | 30.81 | 1.18 |

**Fig. 7.**
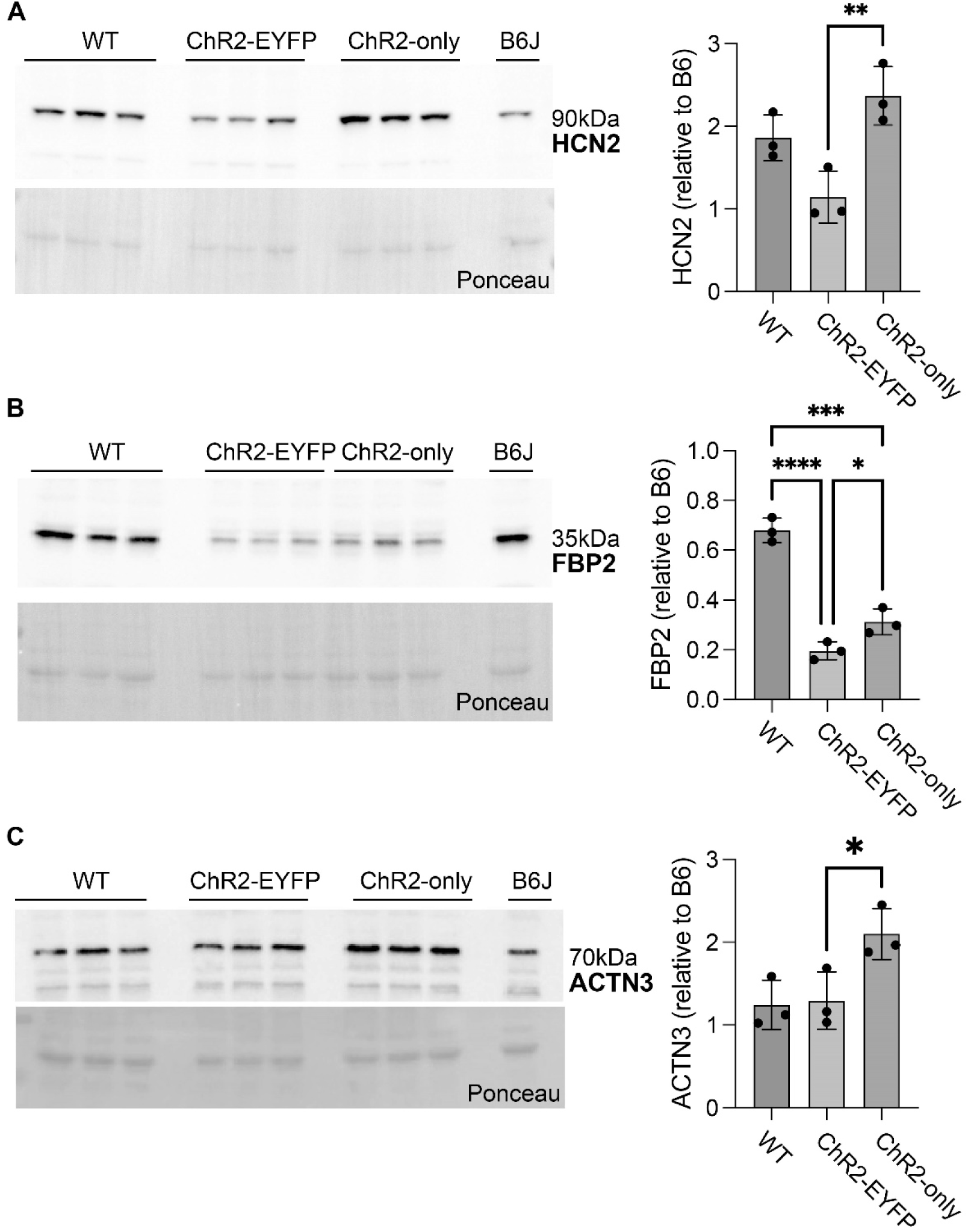
Presence of EYFP led to reduced HCN2 and FBP2 protein levels in gastrocnemius skeletal muscle. Significantly reduced protein-level expressions of (A) HCN2 in the ChR2-EYFP muscle and (B) FBP2 in both ChR2-EYFP and ChR2-only muscles were observed, compared to ChR2-only and WT muscles, respectively. (C) ACTN3 protein expression was significantly higher in ChR2-only muscle compared to ChR2-only muscle. Error bars denote means ± SD. * p<0.05, ** p<0.01, *** p<0.001, and **** p<0.0001.

### Presence of EYFP with ChR2 led to reduced transcriptional sensitivity to optogenetic stimulation compared to ChR2-only muscles

To evaluate if and how muscles from ChR2-EYFP and ChR2-only optogenetic strains differentially respond to daily bouts of repeated optogenetic stimulation, we compared the transcriptomes of stimulated and contralateral unstimulated muscles within each strain. Optogenetic stimulation of ChR2-only muscle resulted in a high number of DEGs compared to the unstimulated side (5,698 DEGs), whereas only 267 DEGs were identified when comparing stimulated ChR2-EYFP muscle to their respective unstimulated side (Table S1). In both strains, we observed an upregulation of inflammatory-response related genes with daily optogenetic stimulation as identified by biological processes enrichment analysis (Fig. 8C and E). In addition to inflammatory response, we also found that ECM organization, collagen organization, and response to mechanical stimulus were enriched by upregulated genes in both strains. Although we observed a small number of biological processes enriched by the downregulated DEGs in stimulated ChR2-EYFP, these all had high enrichment score (Fig. 8B). The downregulated genes in the stimulated ChR2-only genes were primarily associated with mitochondria and metabolism (Fig. 8D).

**Fig. 8.**
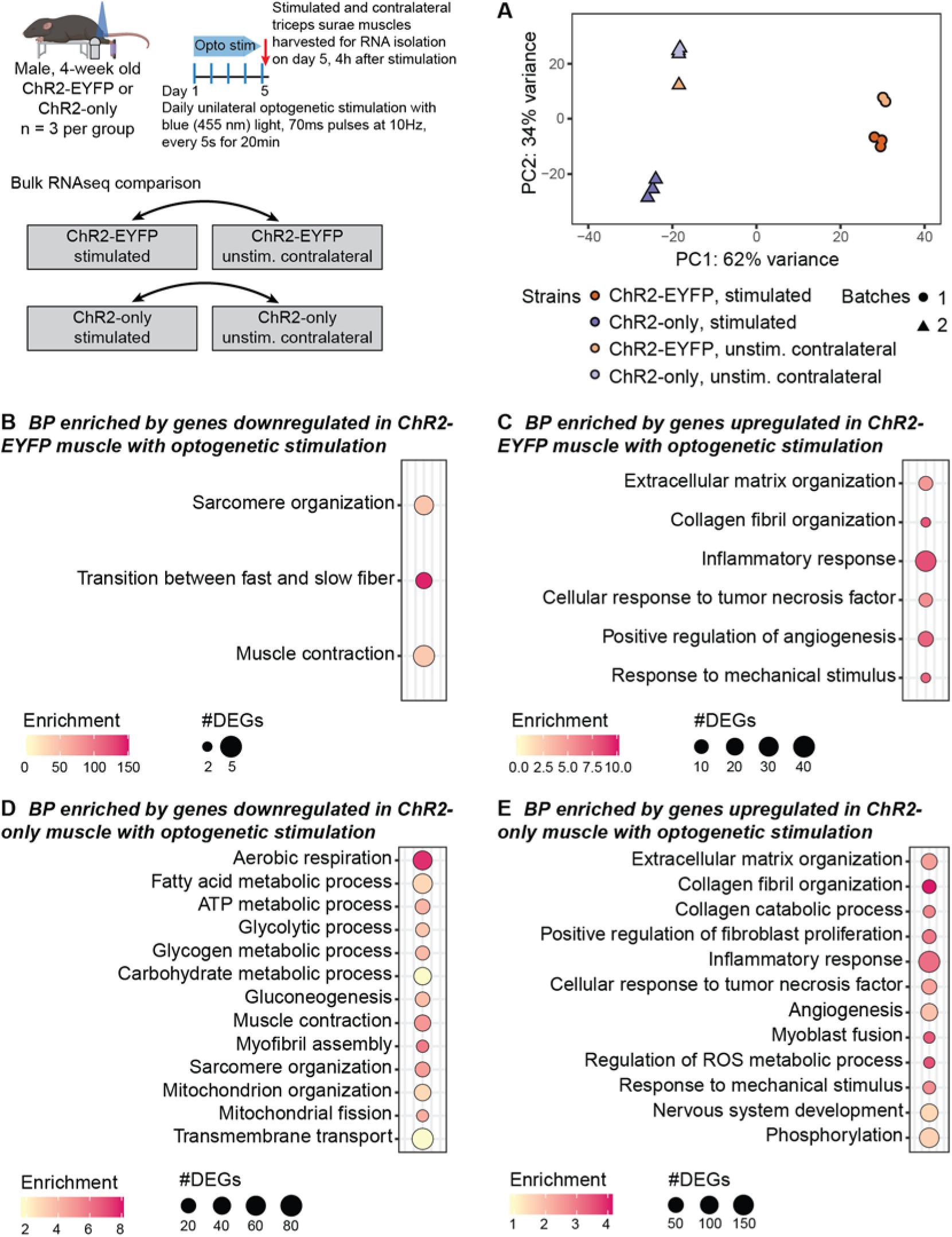
Repeated bouts of optogenetic stimulation led to a more robust transcriptional response in ChR2-only muscle compared to ChR2-EYFP muscle. (A) Principal component analysis of gene expression. Shapes indicate different sequencing batches. Biological processes obtained from genes (D) downregulated and (E) upregulated in ChR2-EYFP stimulated muscle and (E) downregulated and (F) upregulated in ChR2-only stimulated muscle compared to unstimulated contralateral muscle. n = 3 per group.

### In vivo optogenetic stimulation of muscle was less effective primarily due to the presence of mouse skin acting as a barrier to light penetration

To estimate how deep our illumination conditions penetrate the skeletal muscle during optogenetics experiments, we utilized a Monte Carlo (MC) photon transport model. Qualitatively, the Monte Carlo simulations of steady state illumination within the optogenetically simulated tissues reflect expected behavior of photon transport in a turbid biological media (supplementary Fig. S6). In both simulated scenarios, normalized fluence rate (NFR) peaked around the mean free path of the tissue. The mean free path represents the path length a photon will travel on average before changing its direction due to scattering and is equivalent to 1/μs. In the case of the in vitro muscle only results, this is the μs of the muscle, but in the in vivo condition, this is the μs of the most superficial layer, the skin. The photon NFR monotonically decreases deep to this peak in both conditions due to progressive photon energy loss and deposition associated with scattering and absorption events. In the in vivo case, the rate of decay of the decreasing NFR changes at the boundary between the skin and muscle due to the change in underlying optical properties of the medium.

For the single-layer *in vitro* muscle only condition, 23.13% of incident photon energy was absorbed within the simulated tissue cuboid boundaries and 37.6% was absorbed for the two-layer *in vivo* skin-muscle condition (Table 1). For the *in vivo* case, we considered only the total percentage of light that was deposited in the muscle and not the skin. A large percentage of light was reflected in both cases (19.07% muscle only, 30.4% skin-muscle). Despite the highly scattering nature of these tissues, some light (2.94% muscle only, 1.18% skin-muscle) was able to be transmitted through the entire tissue volume. As the x and y edges of the cuboid were simulated as non-reflecting boundaries, any remaining light escaped out the volume from the lateral edges of the cuboid (54.86% muscle only, 30.81% muscle-skin). When considering how much of the incident light penetrated the muscle specifically, this was higher in the muscle only case (23.12%) than the muscle-skin case (8.32%) (Supplementary Table S3). This is due to the increased reflection caused by the skin layer, as well as the fluence that remained in the skin layer itself (4.78%).

In the *in vitro* experiments, an incident irradiance of 140 mW/cm^2^ was used. Therefore, for the cuboid size used in the MC simulations, this would result in an average irradiance of 32.32 mW/cm^2^. Therefore, for the light pulses ranging from 1 ms to 100 ms duration used in these studies, this is equivalent to a 32.32 μJ/cm^2^ to 3.23 mJ/cm^2^ light dose delivered to the muscle volume, respectively. For the *in vivo* experiments, an incident irradiance of 150 mW/cm^2^ was used, resulting in 12.48 mW/cm^2^ average muscle irradiance. This results in a dose of 873.6 μJ/cm^2^ delivered per 70 ms pulse.

### ChR2-EYFP muscles had similar transcriptional profiles as 28-month-old mouse muscles

Given that the ChR2-EYFP muscle exhibited characteristics consistent with aging-associated sarcopenia, including reduced muscle mass, severe force deficit, smaller muscle fibers, and upregulated inflammatory genes (*25–27*), we compared the ChR2-EYFP transcriptome with that of aged mice using SarcoAtlas (https://sarcoatlas.scicore.unibas.ch/) (*28*). SarcoAtlas identified more than 6,000 DEGs in WT aged (28-month-old) mouse muscle compared to young WT (8-month-old) muscle (supplementary Table S2). The overlap of DEGs between the aged and ChR2-EYFP muscle included genes associated with glycolytic processes, gluconeogenesis, and glycogen metabolism, and the overlapping upregulated genes were associated primarily with inflammation.

## Discussion

In this study, we characterized the functional response of optogenetic skeletal muscle with or without the expression of a commonly fused fluorescent reporter, EYFP. Previous studies have explored if and how the overexpression of ChR2-EYFP fusion influences the electrophysiological responses of neuron and HEK293 cells (*17*, *18*). Yet, the role of ChR2-EYFP overexpression in muscle functionality is largely unknown, except for a few studies on the effect of GFP and YFP accumulation in cardiac, neural, and muscle tissues (*12*, *15*, *16*). We developed a new mouse strain by replacing the fluorescent EYFP of light responsive Ai32 with nonfluorescent v5 epitope and expressed this construct in skeletal muscle. Similar to our established ChR2-EYFP mouse model (Acta1-rtTA-tetO-Cre;Ai32) (*6*, *7*), we leveraged this new ChR2-only model to induce optogenetic muscle contraction. We showed that the ChR2-EYFP fusion protein accumulated in the sarcolemma and may also be responsible for abnormal vacuole formation. Expression of ChR2-EYFP led to significant impairments of muscle function, assessed using electrical and optogenetic contractility experiments. With the new ChR2-only mouse model, we overcame this issue and returned muscle contractility using electrical stimulation to that of wildtype mice. These new findings suggest that ChR2-only muscles are physiologically more similar to the WT muscles than the ChR2-EYFP muscles. Green fluorescent protein (GFP) and YFP have been implicated in causing damage in various tissues, including skeletal and cardiac muscles and neurons (*12*). However, if EYFP or ChR2 alone are responsible for functional deficit need further studies. In the new ChR2-only mouse model, ChR2 expression in muscle was reduced compared to ChR2-EYFP muscle, which suggests a potential impairment in translational efficiency as the woodchuck hepatis virus posttranscription regulatory element (WPRE) was removed in addition to EYFP. Thus, the improvements we observed in ChR2-only muscle compared to ChR2-EYFP muscle could be implicated either removal of EYFP or reduced ChR2 expression or both.

Unlike humans, limb muscles in mice are primarily composed of fast-twitch, glycolytic fibers, except for the soleus muscles which consists of around 80% oxidative (type I and IIA) fibers (*29*). Different types of muscles (oxidative and glycolytic) have varied responses to genetic manipulation and stimuli (*30*). In this study, we accounted for different types of muscles, and although we observed significant force and muscle mass deficit in EDL and gastrocnemius muscle, we did not find these same differences in soleus muscles. This differential response may be explained by muscle types, i.e., the ChR2-EYFP overexpression may influence contractility of glycolytic fibers more than oxidative fibers. In line with this differential response based on fiber type, we observed downregulation of fast-twitch muscle-related genes in ChR2-EYFP muscles, however, further studies are needed to better understand this phenomenon. It is also possible that an increase in light irradiance could lead to increased optogenetic-induced contractile force (*4*).

The attenuated muscle force generation in ChR2-EYFP mice that we observed may be in part explained by either decreased calcium handling in muscle cells limited by ChR2-EYFP fusion protein or by structural damage caused by excessive accumulation of EYFP in vacuoles. In our mouse models, ChR2-EYFP co-localized at the sarcolemma and T-tubules (*4*). ChR2-EYFP colocalization may hinder the functionality of voltage gated dihydropyridine (DHP) calcium channels, which could negatively impact the calcium transients, thereby reducing contractile force. We saw a small but significant increase in the half relaxation time of the ChR2-EYFP EDL muscle compared to WT muscle, which may suggest that calcium handling during excitation-contraction coupling was slightly affected by ChR2-EYFP. The attenuated muscle force generation in ChR2-EYFP mice that we observed may also be attributed to potential alterations in the native electrophysiology of muscle fibers caused by ChR2-EYFP overexpression. Elevated ChR2-EYFP can modify cell’s electrophysiological properties (capacitance, thresholds, and currents) (*17*, *18*), which can also decrease conduction velocity of action potentials and inadvertently limit muscle’s force generation capacity and cause contractile dysfunction (*12*, *16*). Additionally, we observed a downregulation in ion-channel associated genes in addition to a reduction in HCN2 protein levels in ChR2-EYFP muscle. Hyperpolarization-activated cyclic nucleotide-gated (HCN) channels activate in response to hyperpolarization and generate an inward cation current that regulates resting membrane potential (*31*). Thus, it is possible that the decrease of HCN2 with ChR2-EYFP overexpression may be impairing the regular electrophysiological activities of muscle fibers. Meng et al demonstrated that expression of ChR2-EYFP in neurons caused slight hyperpolarization of HCN half-activation voltages (*17*), indicating the potential of ChR2-EYFP overexpression to regulate cell’s intrinsic properties. Disrupted calcium handling in muscle fibers induced by ChR2-EYFP fusion protein presents another potential factor contributing to force attenuation. ChR2-EYFP colocalization may hinder the functionality of voltage gated DHP calcium channels; however, additional tests such as calcium transient analysis are required to confirm this. Furthermore, the structural damage caused by excessive vacuole formation in the ChR2-EYFP muscle could also be responsible. Similar membranous whorls are commonly found in muscles exposed to myotoxins such as vincristine, which causes muscle weakness and are hypothesized as a response to the necessity of managing toxic effects (*32*). Thus, it is possible that the functional deficit of muscle is a result of the macro-morphological alterations in muscle cell and tissue structure.

The biological processes enriched by genes downregulated in ChR2-EYFP muscle compared to WT suggest impaired glycolytic metabolism, which may lead to reduced contractile force of EDL and gastrocnemius muscles. This hypothesis is further supported by the downregulation of FBP2 protein which regulates glycogenesis (*33*). The overlap between the downregulated DEGs in 28-month-old mouse muscle obtained from Sarco Atlas and the downregulated DEGs in 3-month-old ChR2-EYFP muscle suggests a severe metabolic dysfunction in the presence of ChR2-EYFP fusion protein. Thus, in addition to functional impairments, metabolic dysfunction in ChR2-EYFP muscle may further compromise the physiological relevance of experimental findings obtained using this optogenetic model. Furthermore, we observed downregulation in fast-twitch fiber contraction related genes which may explain the loss of force in fast-twitch EDL muscle (*21–23*) but not in the slow-twitch soleus muscle. The strong and significant downregulation of *Actn3* is consistent with the functional and structural changes we have observed, as mice lacking *Actn3* have been reported to have reduced muscle strength, mass, and a shift of fast-twitch fibers toward slow twitch fibers (*34*, *35*). The slight increase in half-relaxation time in ChR2-EYFP muscle may also indicate a slowing fiber phenotype, however, additional tests such as single fiber contractility and calcium transient analysis are required to confirm this. We also observed a downregulation of transmembrane transport genes encoding the voltage-gated potassium and sodium ion channels, which may indicate changes in intracellular and transmembrane ion transport mechanisms. Additional contributors of functional impairments could be explained by increased inflammatory responses in the ChR2-EYFP muscle (*36*). The ECM in muscle plays a critical role in contractile force transmission, thus the upregulation of ECM and collagen catabolism related catabolism genes may influence muscle’s force transmission efficiency (*37*, *38*).

Our in vivo 10 Hz optogenetic stimulation protocol generates similar ankle torque as 150 Hz electrical stimulation protocol (*6*). However, ankle torque decays as the muscle gets fatigued and retained at only ∼50% of the initial torque by the end of the 20 min stimulation protocol (data not shown). Generally, an exercise protocol is considered either endurance or resistance type depending on the amount force or torque produced and number of contractions elicited. Skeletal muscle secretes proinflammatory myokines under mechanical loading (*39*), and biological processes indicative of inflammatory and immune responses were upregulated with optogenetic stimulation in both of our optogenetic strains, as previously reported in other studies on exercise (*40*, *41*). However, the ChR2-only muscle was more sensitive than the ChR2-EYFP muscle as the same stimulation protocol yielded a higher number of DEGs in the ChR2-only muscle. We observed downregulation of genes related to metabolism and mitochondrial regulation in the stimulated ChR2-EYFP muscle; however, this phenomenon is not uncommon in resistance exercise (*42*). Viggars et al demonstrated that with daily electrical stimulation in rats, mitochondrial genes were downregulated in rat skeletal muscle until training period reached 30 days (*41*). Thus, our optogenetic muscle stimulation protocol may be considered resistance exercise as it produces high torque with fewer contractions (*43*, *44*).

This study holds great significance in the field of musculoskeletal bioengineering as we demonstrated the potential adverse effects of fluorescent Cre reporters on muscle physiology. Cre reporter strains are popular in biomedical research for a variety of tasks, such as cell lineage tracing, validating recombination, tracking cellular processes, and monitoring expression in living tissues and cultured cells. However, our findings revealed significant dysfunctions in skeletal muscle expressing ChR2 with EYFP (i.e., Ai32). Upon EYFP removal (ChR2-only), we observed substantial improvement in muscle contractility and physiological response to stimulation. This work has notable implications for the field, as it shows the potential to improve ChR2 delivery and efficacy in skeletal muscle thus strengthening the translational potential of optogenetics for regulating muscle function.

## Methods

The Unit for Laboratory Animal Medicine at the University of Michigan approved all animal procedures.

### Animal models

All animals were kept on a 12-hour light/ dark cycle under standard housing conditions with ad libitum access to water and chow throughout the experiment. Mice were genotyped by PCR (Transnetyx, TN, USA). Wildtype (WT) littermates were used as controls. Acta1Cre;ChR2-EYFP mice were generated as described previously (*6*). Briefly, tetracycline inducible Acta1Cre (*45*) male mice were bred with Ai32^fl/fl^ (Jax ID: 024109, C57BL6/J background) female mice to generate Acta1Cre;Ai32^fl/fl^ mice, referred to as ChR2-EYFP onward. Cre recombination was induced via doxycycline (tetracycline) chow (Inotiv, 200 Doxycycline) at mating and continued till pups were weaned. The ChR2-only mouse line was generated by deleting the EYFP-WPRE DNA fragment from the ChR2-EYFP mice (Jax Stock No. 024109) using the CRISPR/Cas9 technology (*46*). Briefly, two single guide RNAs (sgRNA), one targeting the junction between ChR2 and EYFP (AGTACCCGCGGCCGCCACCA) and the other one targeting the end of WPRE (GGCCCTAGGCGGTATCGATG in reverse orientation), were generated using ThermoFisher’s in vitro transcription service. These two sgRNAs (20 ng/ul each) were co-microinjected with Cas9 mRNA (50 ng/ul, TriLink BioTechnologies) into the cytoplasm of zygotes collected from ChR2-EYFP mouse mating pairs. A single-strand oligonucleotide donor (GAGGTCGAGACGCTGGTGGAGGACGAGGCCGAGGCTGGCGCAGTACCCGCGGCCGCCA CC**GGTAAGCCTATCCCTAACCCTCTCCTCGGTCTCGATTCTACG**CGATACCGCCTAGGGCC TCGACTGTGCCTTCTAGTTGCCAGCCATCTGTTGTTTGCCCCT) was also included in the injection mix (100ng/ul) for mediating precise deletion as well as adding a V5 tag (sequence in bold) to the C-terminus of the ChR2 transgene. Injected embryos were cultured overnight in KSOM medium (Millipore Sigma) in a 37°C incubator with 6% CO_2_. In the next day, embryos that had reached the 2-cell stage of development were implanted into the oviducts of pseudo-pregnant surrogate mothers (CD1 strain from Charles River Laboratory). Offspring born to the foster mothers were genotyped using PCR followed by Sanger sequencing. Founder mice with the desired deletion V5 tagging were expanded to establish the ChR2-only mouse line. Acta1Cre;ChR2-only^fl/fl^ mice, referred to as ChR2-only onward were generated similarly as ChR2-EYFP mice. For light and electron microscopy, constitutive CKCre;Ai32^fl/fl^ mice were also used and were generated in Glancy laboratory.

### Light and electron microscopy

Extensor digitorum longus (EDL) muscles from adult (> 2-months old) WT and ChR2-EYFP male mice were flash-frozen for histology following standard protocols (n=1/group). Tissues were sectioned at 10 um thickness longitudinally or transversely and were imaged in confocal or widefield microscope. For transmission electron microscopy (TEM), EDL muscles from adult WT, ChR2-EYFP, and ChR2-only male mice were fixed with glutaraldehyde and osmium tetroxide and were embedded in plastic (n=1/group). Ultrathin sections were produced to visualize the ultrastructure in JEOL-1400 transmission electron microscope.

### In situ contractility of gastrocnemius muscle using nerve stimulation

Experiment was conducted as detailed in Larkin et al 2011 (*47*). Briefly, 3-month-old WT, ChR2-EYFP, and ChR2-only male (n = 4, 3, and 5, respectively, per group) mice were anesthetized using 2,2,2-Tribromoethanol suspended in 2-methyl-2-butanol and sterile saline. 0.3-0.5 mL anesthetic was injected intraperitoneally as frequently as needed to maintain anesthesia. The mouse under anesthesia was placed on a heated platform to maintain optimal body temperature throughout the procedure. The gastrocnemius muscle was carefully separated from the surrounding musculature without damaging the blood supplies and nerves. A suture was tied around the Achilles tendon and calcaneus bone and then the calcaneus was dissected. The knee and foot were tied to fixed posts to limit motion and maintain isometric condition. The Achilles tendon was tied to a dual mode lever system (6650LR, Cambridge Technology). The sciatic nerve under the hamstring muscle was exposed and a bipolar platinum wire electrode was placed upon it. Heated saline (37°C) was dripped continuously on the muscle to prevent it from drying. The gastrocnemius muscle was activated by stimulating the nerve with electrical pulses to induce contraction. Stimulation voltage and muscle length were adjusted to obtain maximum twitch force. Trains of electrical pulses were then applied to produce isometric tetanic contractions. Forces at frequencies ranging from 40-220 Hz were recorded. We normalized the maximum isometric tetanic force with respect to the physiological cross-sectional area (PCSA) of the muscle using equation (3.1)

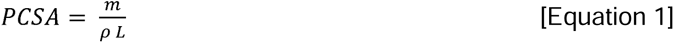

Where, m = muscle mass, ρ = density (1.06 g/cm^3^), and L = muscle fiber length.

### In vitro contractility of EDL and soleus muscles

#### Electrical stimulation

Experiment was conducted as detailed in Brooks and Faulkner 1988 (*48*). Briefly, EDL muscles were dissected from 3-month-old WT, ChR2-EYFP, and ChR2-only male mice (n = 5, 3, and 5, respectively, per group) under anesthesia. The isolated EDL muscle was placed in buffered mammalian Ringer solution bath to maintain physiological condition. Distal and proximal tendons were tied with sutures to a fixed post and force transducer (BG-50, Kulite semiconductor), respectively. Two platinum electrodes were placed on either side of the muscle parallel to its long axis. An electrical field was applied between the electrodes and the muscle was tested following the same protocol as the in situ test described before. Time to peak twitch force and half relaxation times were also recorded. Soleus muscles were tested similarly following this same method (n = 3, 3, and 4, respectively, per group, all male except two in the ChR2-EYFP group). Maximum isometric tetanic forces were normalized with respect to respective muscles’ PCSA using equation [1].

#### Optogenetic stimulation

With the muscle remaining in the same experimental chamber as was used for electrical stimulation, we optogenetically stimulated EDL (n = 3 and 4, respectively, per group) and soleus (n = 3 and 4, respectively, per group, all male except two in the ChR2-EYFP group) muscles following electrical stimulation. A blue light (model, 455 nm wavelength, Thorlabs) was placed directly above the muscle at an irradiance of 140 mW/cm^2^. We measured muscle peak twitch force with incremental light pulse duration ranging from 1 to 100 ms. The ability of the muscle to sustain tetanic contraction with optogenetic stimulation was also tested by varying LED pulse durations and frequency over 300 ms duration.

### Muscle fiber cross-sectional area (CSA) measurements

The fiber cross-sectional area (CSA) was determined using the contralateral unstimulated EDL muscles from the mice utilized in the contractility experiments (n = 3 per group). The EDL muscle was flash-frozen following standard histology protocols. 10 um thick transverse sections were obtained from the mid belly region and were stained with Alexa 647 fluorophore conjugated wheat germ agglutinin (WGA). Sections were imaged in fluorescence microscope (BioTek Lionheart FX with Cy5 cube) and fiber CSA were determined with FIJI particle analysis.

### RNA sequencing of muscles

#### In vivo muscle stimulation

1-month-old ChR2-EYFP and ChR2-only mice (n = 3 per genotype) were subjected to isometric unilateral optogenetic muscle stimulation for five days, as described previously (*7*). Briefly, with the mouse in prone position, right limb was placed in an in-vivo muscle stimulation apparatus (Aurora Scientific, ON, Canada), with the foot resting against a flat plate. The limb was clamped at the knee to stabilize the joint for isometric triceps surae contractions. A custom-built setup was used to shine controlled light pulses on the tricep surae (*6*), which consisted of a blue-light LED (455 nm, 900 mW, Thorlabs, NJ, USA) and a driver (DC2200, Thorlabs) for pulse modulation. Muscle was stimulated at 10 Hz frequency (70 ms on-/30 ms off-time; 10 repetitions/ second) for 20 minutes for five days. The left limbs served as unstimulated internal controls. After each daily session, mice were returned to cages to resume regular activity.

#### RNA isolation and sequencing

Mice were euthanized following institutional guidelines four hours after their last stimulation session. Age matched WT mice were euthanized at the 5-day time points. From stimulated mice, tricep surae muscles of both limbs were rapidly dissected, flash-frozen in liquid nitrogen, and stored in −80 C until RNA isolation. From WT mice only the right tricep surae muscles were harvested. Samples were pulverized in TRIzol (Ambion, Austin, TX, USA) using Precellys tissue pulverizer. total RNA was extracted using Invitrogen PureLink RNA Mini Kit with on-column DNA digestion (Ambion). RNA was eluted in 50 uL RNase free water. RNA integrity number (RIN) was measured using Agilent bioanalyzer. A total of 15 muscle samples were sequenced in two different batches. Next generation sequencing was performed on poly-A mRNA libraries as described before (*7*).

### Computational estimation of light penetrance during optogenetic stimulation

We utilized a Monte Carlo (MC) photon transport model to quantify light penetration depth in muscle tissue for our optogenetics experiments (*49*, *50*). MC simulations are physics-based, stochastic models of photon propagation through a turbid biological medium. Individual photons are launched into a simulated tissue with user-specified optical properties (scattering coefficient [µ_s_ (cm^-1^)], absorption coefficient, [µ_a_ (cm^-1^)], scattering anisotropy factor [*g*]). Photon movement and local energy deposition is tracked until the photon has either lost all its original energy or exited the tissue volume (supplementary Fig. S6). This process is repeated for many photons (∼10^6^-10^8^) such that average distribution of the light propagation path within the tissue can be reliably estimated. By tracking the locally deposited power of each photon during the simulation, the normalized fluence rate, or NFR, (cm^-2^) can be output for each voxel within the tissue cuboid. This conceptually represents the fraction of irradiance that was incident on that voxel during the simulation per Watt of incident light.

To perform these simulations, an open source, MATLAB based MC model (“MC Matlab”) was used. A one- or two-layer model was generated to describe both the *in vitro* (muscle only) and *in vivo* (skin + skeletal muscle) experiments, respectively. Following optical properties were extracted from relevant literature: scattering coefficient, µ_s_ = 200 and 90 cm^-1^, absorption coefficient, µ_a_ = 6 and 1 cm^-1^, scattering anisotropy factor, *g* = 0.75 and 0.9, for skin and muscle, respectively (*51–54*). Refractive indices were assumed 1.4 for both tissues. The light source was described as an infinite plane wave at 455 nm with normal incidence on the surface of the tissue. For each simulation, a 3D volume of 100 x 100 x 300 bins was generated for a total tissue volume of .35 x .35 x .35 cm (voxel size of .035 x .035 x .012 mm; length, width, height, respectively). A total of 10^8^ photons were launched per simulation. All cuboid boundaries were set as non-reflective to best mimic native conditions and not forcibly and erroneously “trap” light within the simulated sub volume of the muscle. As such photons could escape from the lateral edges of the volume and not be “reflected”, “transmitted”, or “dead” within the tissue volume.

### Western blotting

Following euthanasia, gastrocnemius and soleus muscles from 5-week-old mice (n = 3 per group) were dissected, flash-frozen in liquid nitrogen, and stored in −80 C until protein extraction. Briefly, 20-30 mg of tissue was lysed in radio immunoprecipitation assay (RIPA) buffer supplemented with protease and phosphatase inhibitors. Following lysis, the suspension was centrifuged, and the resulting supernatant was collected for blotting. Protein concentration was measured with Pierce BCA protein assay (Thermo Scientific, #23225). For ChR2 blotting, 10 μg gastrocnemius muscle protein diluted in loading buffer was reduced with dithiothreitol and was loaded onto 8-16% tris glycine gel. Following running, proteins were transferred to polyvinylidene difluoride (PVDF) membrane. The membrane with transferred protein was stained with Ponceau S stain (Sigma-Aldrich #P7170) to check the quality of transfer and protein load. The membrane was incubated with anti-ChR2 primary antibody (Table 2). A separate membrane was developed with soleus muscle protein similarly. Other targets, TIMP1, FBP2, ACTN3, and HCN2 were selected from the sequencing result based on their differential expression intensity. For these targets, 15 μg gastrocnemius muscle protein was denatured and reduced before loading onto the gel and primary antibodies were used at dilution ratios mentioned in Table 2. The membranes were then incubated with horseradish peroxidase-conjugated secondary antibody (Table 2). Protein bands were developed with enhanced chemiluminescent substrate (SuperSignal™ West Pico PLUS Chemiluminescent Substrate, #34577) and were imaged with ChemiDoc MP imaging system (Bio-Rad). All blots included a well with protein isolated from the gastrocnemius muscle of a 6-month-old C57BL6/6J female mouse. Band intensity for immunoblots were normalized with respect to representative bands from total protein staining (Ponceau S) or to intensity of the C57BL6/J band. Images were analyzed with FIJI Gel plugin.

**Table 2:**
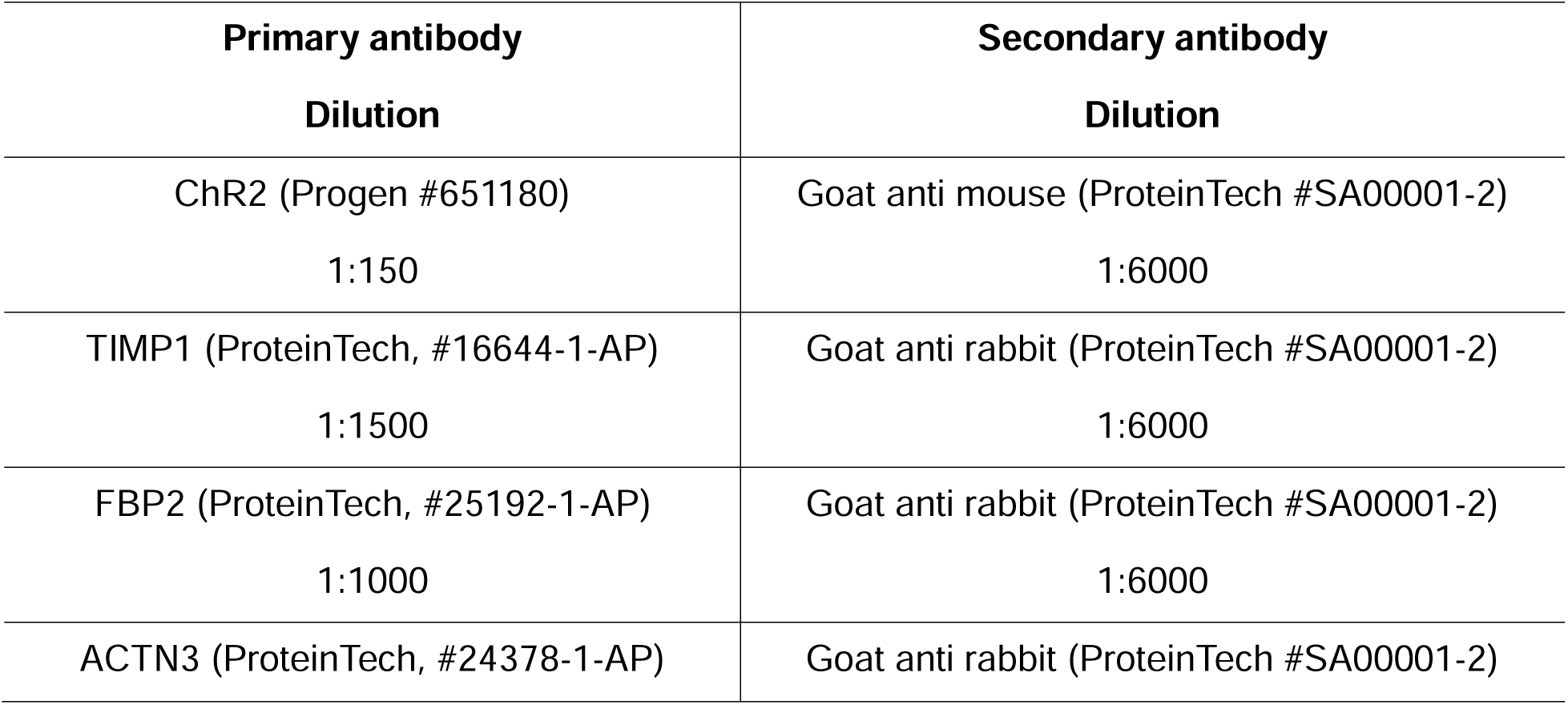
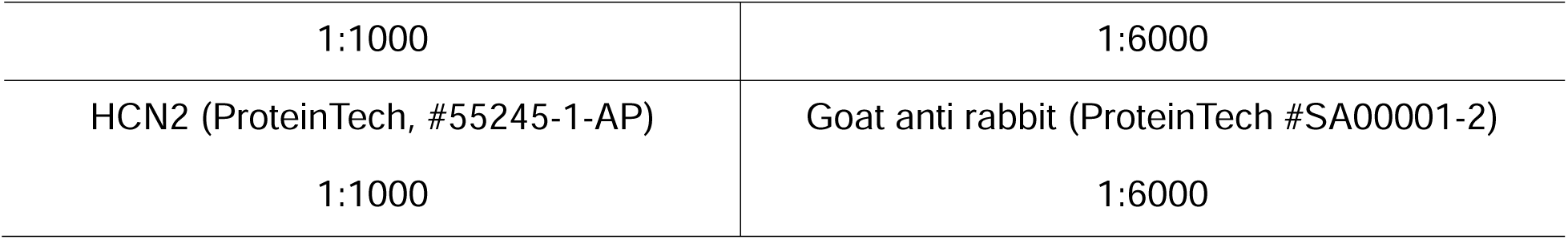
Western blot antibody information.

### Statistical analysis

Statistical analyses were performed using GraphPad Prism (version 9 or higher, La Jolla, USA) or in RStudio (version 4.1.0).

#### Contractility, muscle fiber CSA, and western blot measurements

Body masses, muscle masses, maximum specific tetanic forces, and muscle fiber CSAs were analyzed with one way ANOVA corrected for multiple testing (Tukey’s post-hoc) across genotypes. Tetanic forces were compared using two-way repeated measures ANOVA to determine significance across genotypes and stimulation frequencies. Significance was set at p<0.05. For western blots, band intensities (calculated as adjusted volumes in ImageLab software) were compared using one-way ANOVA with Sidak’s correction for multiple comparisons.

#### Differential expression of genes (DEG)

We used DESeq2 in R/Bioconductor (*55*) to determine the DEGs. Count data were organized in two different count matrices: matrix 1: WT, ChR2-EYFP unstimulated, and ChR2-only unstimulated; and matrix 2: ChR2-EYFP unstimulated, ChR2-EYFP stimulated, ChR2-only unstimulated, and ChR2-only stimulated. In matrix 1, we analyzed ChR2-EYFP and ChR2-only unstimulated samples with WT as baseline controlling for batch variability. In matrix 2, ChR2-EYFP and ChR2-only stimulated samples analyzed with their respective unstimulated samples as baseline controlling for batch, genotype, and animal variability. Principal component analysis (PCA) was performed with the top 500 most variable genes across samples in a matrix to visualize sample clustering. For analyses in matrix 1, significance was tested against log_2_(FoldChange) = 0.15, while in matrix 2 significance was tested against 0.32. p-values were adjusted for multiple testing using the Benjamini and Hochberg method, and differential expression significance was set at p-adjusted (p-adj) <0.05.

To compare our dataset with aged muscle, we used SarcoAtlas (https://sarcoatlas.scicore.unibas.ch/). Differential expression analysis was compared using the mouse gastrocnemius time course data (*28*). We determined DEGs between 28-month-old and 8-month-old mouse muscle with 8-month-old muscle as baseline. A threshold of log2(FoldChange) = 0.15 was applied and significance was set at FDR < 0.01.

#### Enrichment analysis

To determine enriched biological processes, the list of up and downregulated DEGs were input separately into the database for annotation, visualization, and integrated discovery (DAVID) (*56*, *57*). A threshold of false discovery rate (FDR) <0.1 was applied to identify significantly enriched gene ontology biological processes.

