## Supplemental for "Overexpression of enhanced yellow fluorescent protein fused with Channelrhodopsin-2 causes contractile dysfunction in skeletal muscle"

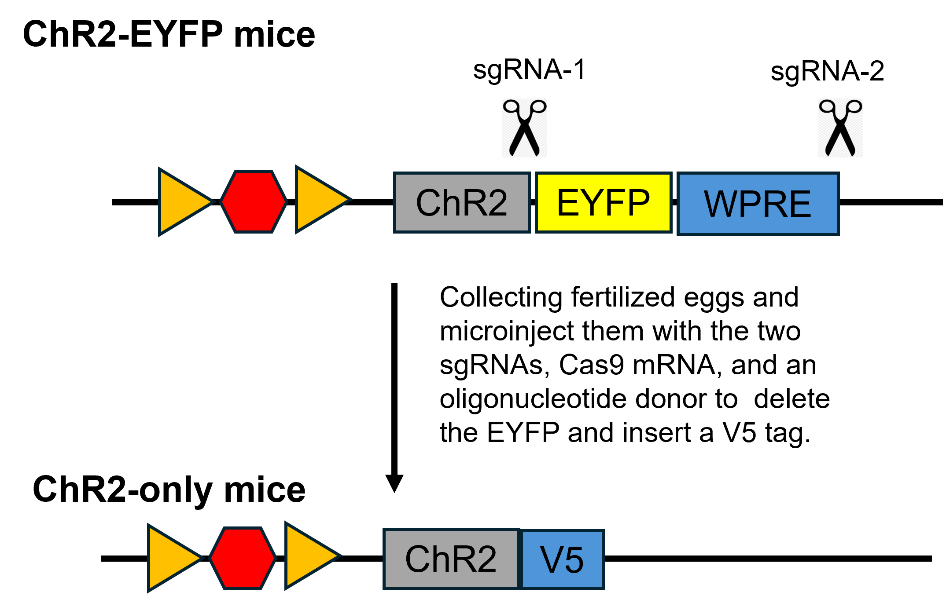

**Fig. S1. Generation of ChR2-only mouse.** EYFP-WPRE DNA fragment from the ChR2-EYFP mice were deleted from the ChR2-EYFP mice using the CRISPR/Cas9 technology.

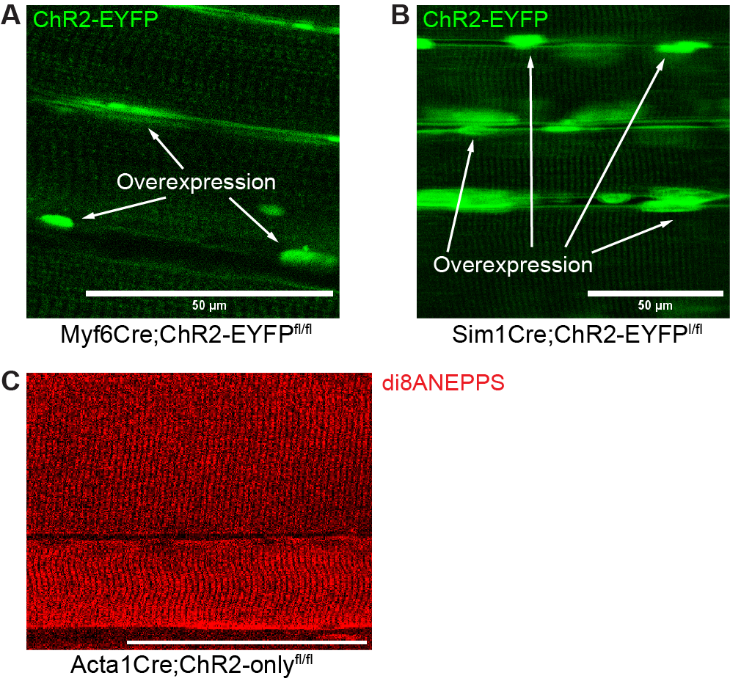

**Fig. S2. Abnormal ChR2-EYFP clustering in skeletal muscle was observed in different optogenetic mouse models.** Confocal imaging demonstrated ChR2-EYFP overexpression in the EDL muscle of (A) Myf6Cre;ChR2-EYFP^fl/fl^, (B) CKCre;ChR2-EYFP^fl/fl^ muscle. (C) The di-8-ANEPPS-stained EDL muscle of the Acta1Cre;ChR2^fl/fl^ mouse showed a regular muscle cell membrane and T-tubule structure.

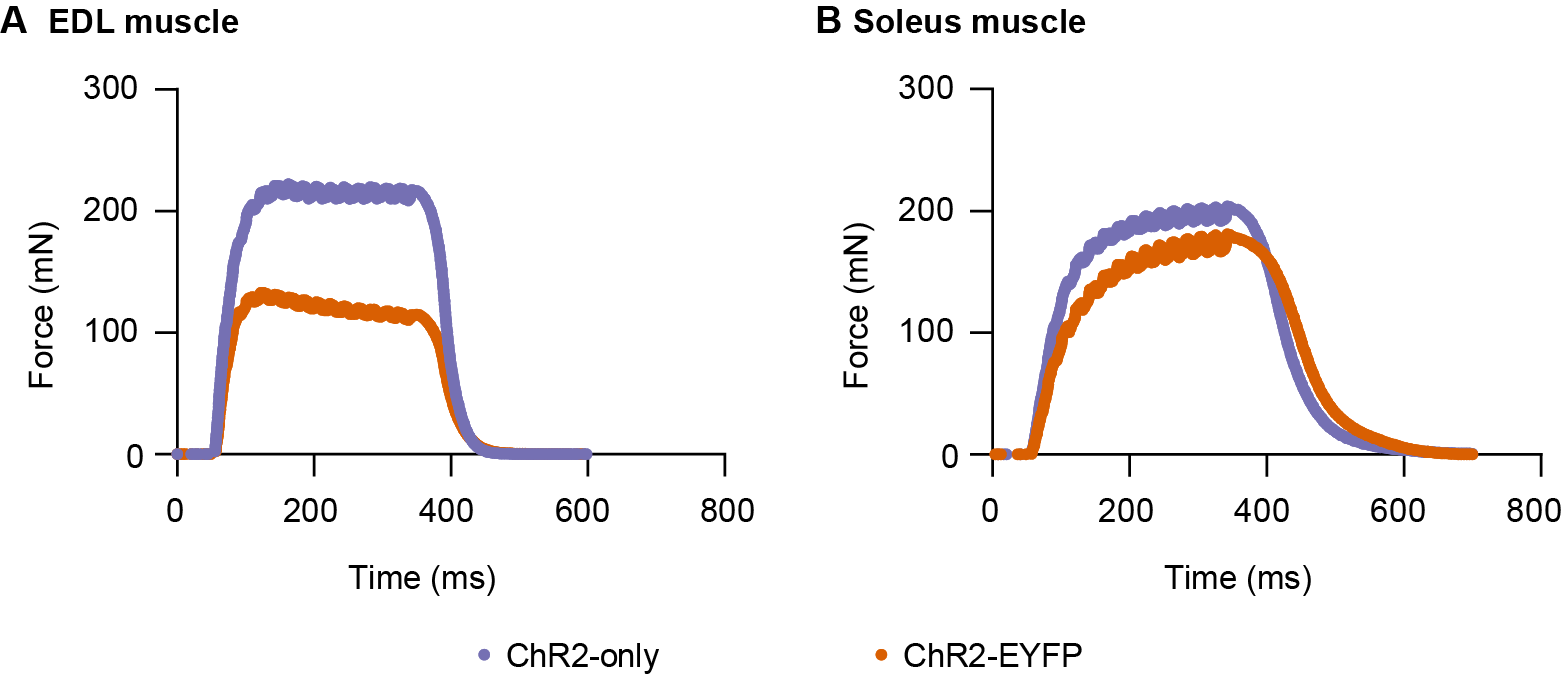

**Fig. S3. Optogenetic stimulation of muscle elicits partially fused tetanic contraction in vitro.** Representative tetanus at 50 Hz, 10 ms pulse duration in (A) EDL and (B) soleus muscle.

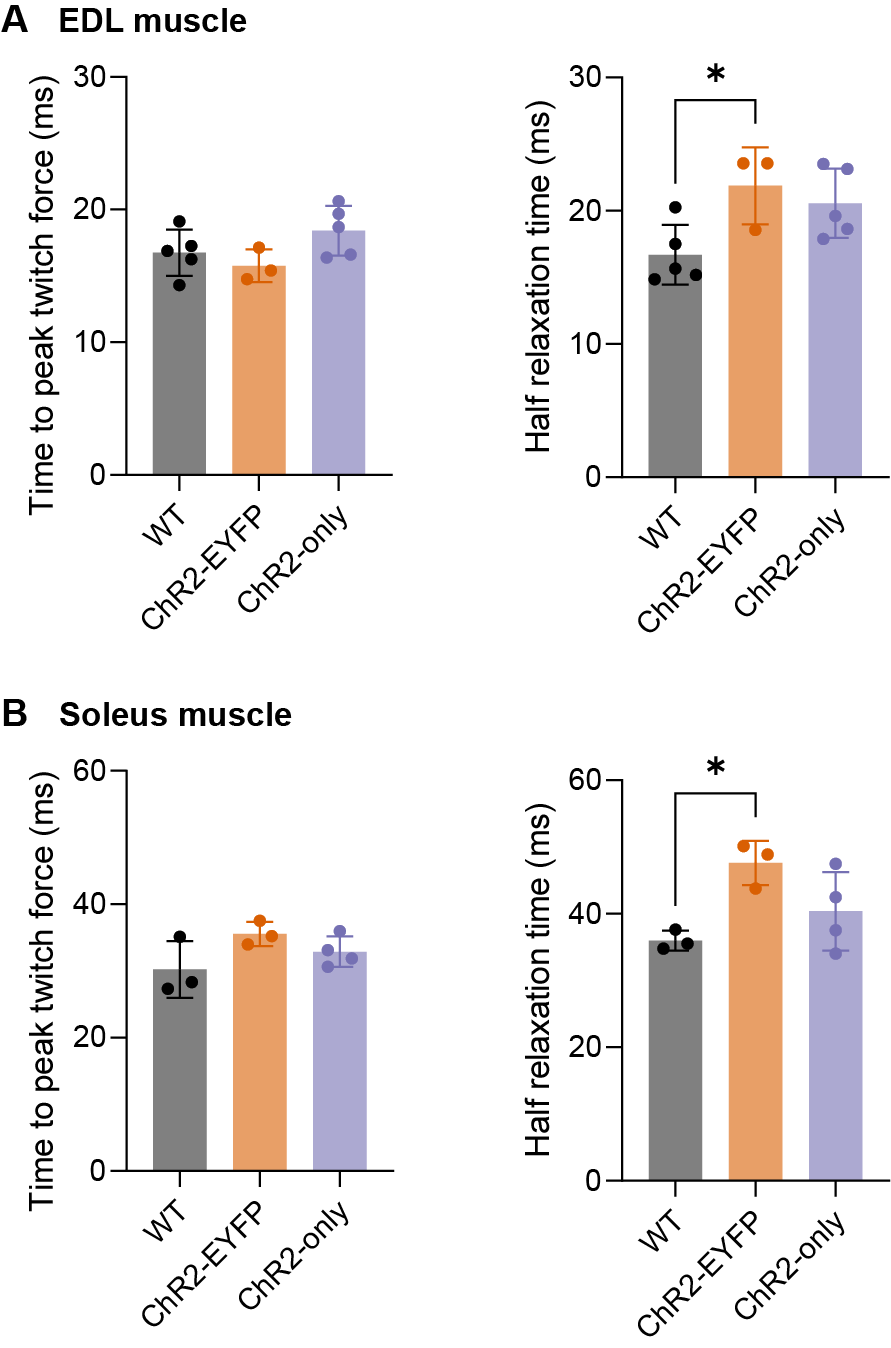

**Fig. S4. ChR2-EYFP overexpression in muscle led to increased muscle half relaxation time.** Twitch times of (A) EDL and (B) soleus muscles**.** Data were compared using one-way ANOVA with Tukey’s correction for multiple tests. Error bars denote means ± SD. * p<0.05.

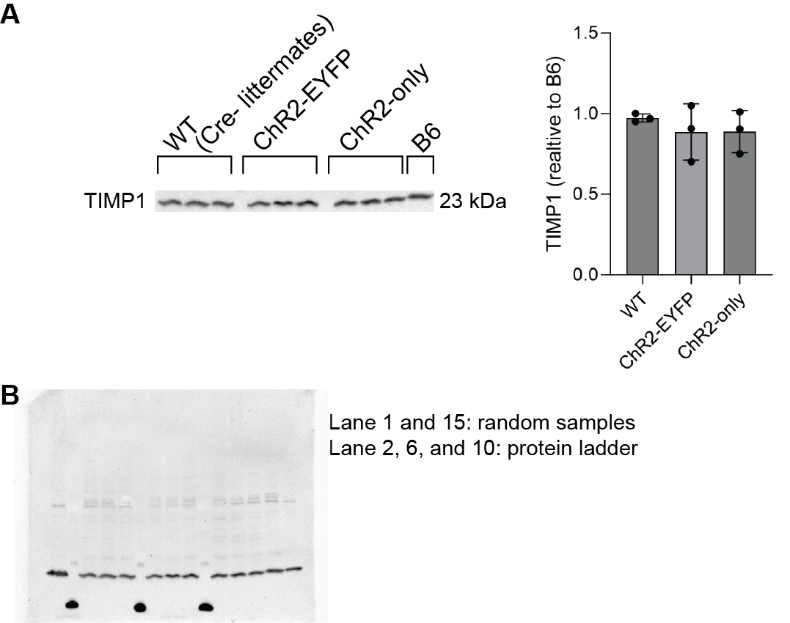

**Fig. S5. Western blot analysis of TIMP1 in skeletal muscles.** (A) TIMP1 expression was not different across WT, ChR2-EYFP, and ChR2-only gastrocnemius muscles. Band intensity for immunoblots were normalized with the intensity of the C57Bl6/J band. (B) TIMP1 whole blot image.

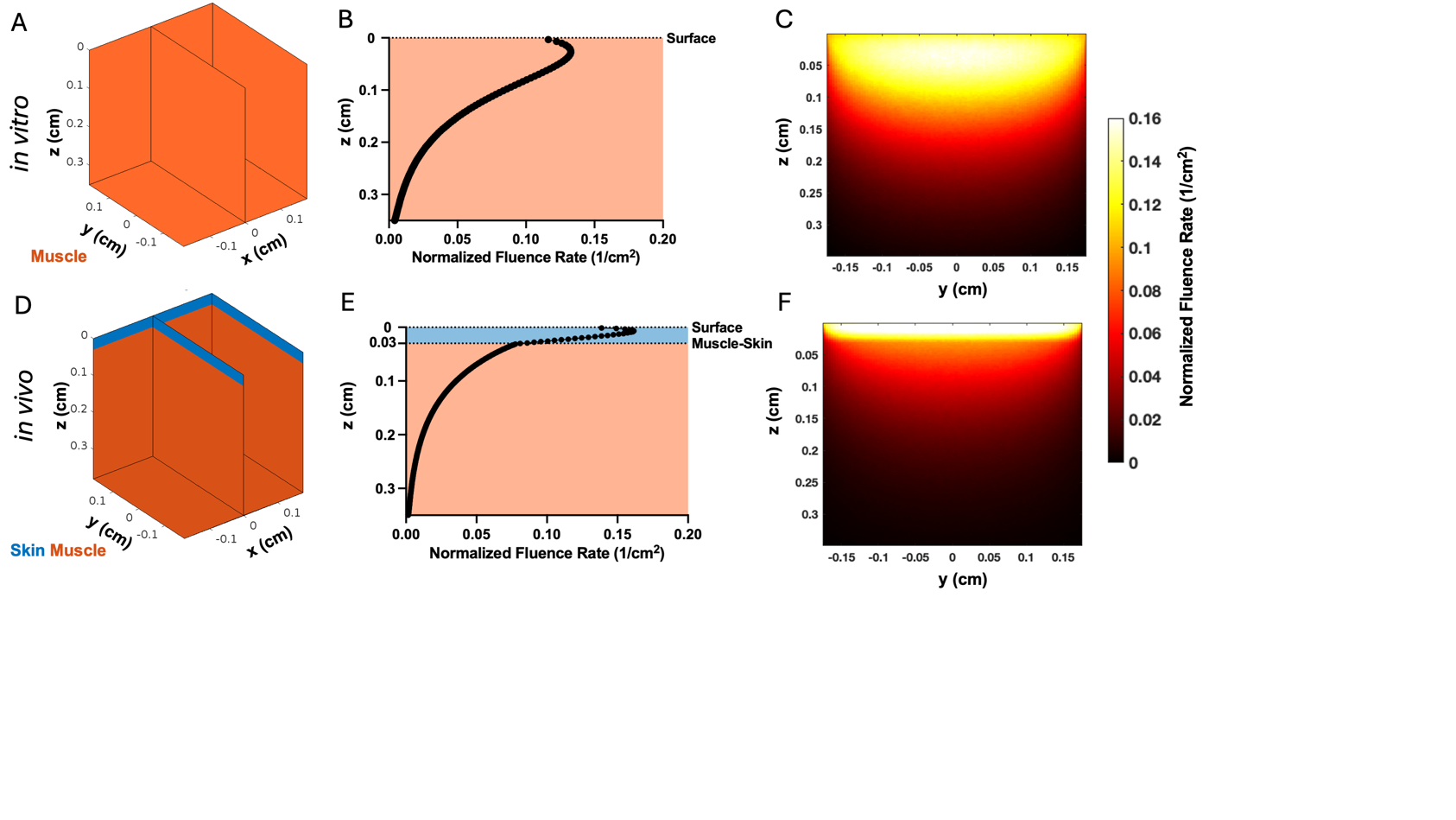

**Fig. S6.** Monte Carlo photon transport simulations of light penetrance in (**A-C**) *in vitro* and **(D-F**) *in vivo* optogenetic stimulation experiments. (**A, D**) Geometrical representation of simulated tissue consisting of (**A**) muscle only, or (**D**) skin + muscle. (**B,E**) Normalized fluence rate (NFR [cm^-2^]) as a function of depth. **(C,F**) NFR across the 2D plane of the tissue volume defined in (**A, D**) where x = 0.

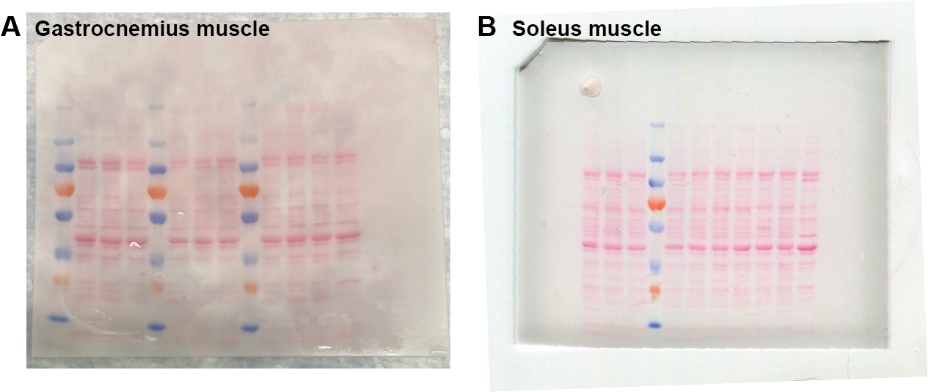

**Fig. S7. Ponceau stain of ChR2 blots shown in Fig. 3.** Equal stain intensity was observed in all lanes along the same molecular weight.

**Table S1. Numbers of differentially expressed genes (DEGs) in triceps surae muscles in RNA-seq experiments.** Optogenetically stimulated muscles were harvested 4 h after last bout.

| **RNA-seq comparisons** | **Total #DEGs** | **#DEGs downregulated compared to baseline** | **#DEGs upregulated compared to baseline** |
| --- | --- | --- | --- |
| ChR2-EYFP unstimulated vs WT | 1,283 | 632 | 651 |
| ChR2-only unstimulated vs WT | 410 | 251 | 159 |
| ChR2-EYFP stimulated vs unstimulated contralateral | 267 | 47 | 220 |
| ChR2-only stimulated vs unstimulated contralateral | 5,698 | 2,546 | 3,152 |

**Table S2: Differential expression from SarcoAtlas RNAseq.** Comparision was made between 28-month-old and 8 month old mouse muscles

| **SarcoAtlas RNA-seq comparison** | **Total #DEGs** | **#DEGs downregulated compared to baseline** | | **#DEGs upregulated compared to baseline** |
| --- | --- | --- | --- | --- |
| 28-month vs 8-month | 6, 229 | 3,175 | | 3,454 |
|  | **Downregulated** | | **Upregulated** | |
| **#DEG overlap with ChR2-EYFP** | 221 | | 170 | |
| **Biological processes enrich by overlap (enrichment)** | Glycolytic process (15.3), gluconeogenesis (13.1), glycogen metabolic process (13.4) | | Inflammatory response (5.5) | |

**Table S3. Percentages of total incident light fluence relative to locations within cuboid.**

|  | % of incident light fluence in skin | % of incident light fluence in muscle | % of incident light fluence in most superficial 1mm of muscle | % of incident light fluence in most superficial half of muscle |
| --- | --- | --- | --- | --- |
| in vitro | - | 23.13 | 14.22 | 19.55 |
| In vivo | 4.78 | 8.32 | 5.80 | 7.37 |
